# sCCIgen: A high-fidelity spatially resolved transcriptomics data simulator for cell-cell interaction studies

**DOI:** 10.1101/2025.01.07.631830

**Authors:** Xiaoyu Song, Joselyn C. Chavez-Fuentes, Weiping Ma, Weijia Fu, Pei Wang, Guo-Cheng Yuan

## Abstract

Spatially resolved transcriptomics (SRT) provides an invaluable avenue for examining cell-cell interactions within native tissue environments. The development and evaluation of analytical tools for SRT data necessitate tools for generating synthetic datasets with known ground truth of cell-cell interaction induced features. To address this gap, we introduce sCCIgen, a novel real-data-based simulator tailored to generate high-fidelity SRT data with a focus on cell-cell interactions. sCCIgen preserves transcriptomic and spatial characteristics in SRT data, while comprehensively models various cell-cell interaction features, including cell colocalization, spatial dependence among gene expressions, and gene-gene interactions between nearby cells. We implemented sCCIgen as an interactive, easy-to-use, realistic, reproducible, and well-documented tool for studying cellular interactions and spatial biology.

## Introduction

Understanding cell-cell interactions is essential for unraveling the complexities of biological systems and advancing medical science. Cell-cell interactions guide cells to form tissues and organs in a precise and coordinated manner. In the immune system, their interactions enable the efficient detection and response to pathogens^1^. Aberrant interactions are often at the heart of diseases. For example, disrupted communication between neurons leads to cognitive decline and memory loss in Alzheimer’s disease^2^, while abnormal interactions among cancerous and immune cells facilitate tumor growth and metastasis^3^. Treatments targeting these cellular interactions, such as the PD-1/PDL-1 antibodies targeting the tumor immune escape^4^, have demonstrated remarkable effectiveness in clinical settings. Therefore, studying cell-cell interactions is fundamental not only for understanding biological intricacies but also for driving innovations in health and medicine.

Spatially resolved transcriptomics (SRT) technologies serve as an invaluable tool for exploring cell-cell interactions, by enabling researchers to examine the gene expression profiles of cells within the native tissue context^5,6^. By preserving spatial information, SRT facilitates the study of how the physical distribution of cells relates to their interactions. Various computational tools have been developed to analyze these interactions: some focus on analyzing the spatial colocalization of cells on tissue slides^7^; others explore the association between gene expression levels and the cell-to-cell distances (expression-distance association)^8,9^; and some examine the co-expression of genes, such as from ligand-receptor pairs, in nearby cells (expression-expression association)^10–14^.

Understanding the performance of these rapidly evolving SRT technologies and computational tools is critical for their effective applications. However, this task is challenging due to the lack of a standardized reference dataset with known ground truth. Simulations can help, but existing SRT data simulators, such as scDesign3^15^ and SRTsim^16^, have limitations in capturing intricate cell-cell interactions (Table 1). scDesign3^15^, extended from a scRNAseq data simulator, excels in capturing transcriptomics features, such as gene-gene correlations and sequencing depth, while SRTsim^16^ specializes in capturing spatial features, including accommodating different spatial inputs and determining spatial windows to house the simulated cells. Although both simulators preserve some spatial expression patterns, neither effectively captures cell-cell interactions in generating spatial and transcriptomic data. Additionally, given the scarcity of SRT data compared to the relatively abundant scRNAseq/snRNAseq data, especially for hard-to-obtain tissues and conditions, neither simulator can leverage single-cell transcriptomics to simulate the SRT data. Finally, both simulators focus on reproducing the existing SRT patterns when reference data is given, falling short in generating *de novo* patterns for evaluations under diverse conditions. Evaluating beyond the limited patterns captured in existing data is crucial, as tissues, such as tumors, can be heterogeneous and exhibit distinct spatial patterns. Therefore, there is a pressing need to develop an additional SRT data simulator that incorporates these nuanced patterns. Such a simulator would significantly advance the development and testing of SRT computational tools and analysis pipelines.

**Table 1:** The comparison between sCCIgen and the existing SRT simulators. sCCIgen offers real-data-based simulations with a strong emphasis on cell-cell interactions. It not only faithfully reproduces both spatial and expression patterns observed in real datasets, but also has the capability to generate *de novo* patterns to address data availability limitations.

|  |  | scDesign3 | SRTsim | sCClgen (Proposed) |
| --- | --- | --- | --- | --- |
| Input | SRT | ✓ | ✓ | ✓ |
|  | No data |  | ✓ |  |
|  | Single-cell RNA seq |  |  | ✓ |
| Spatial | Estimate window |  | ✓ | ✓ |
|  | Mimic real cell allocation |  |  | ✓ |
|  | **Allow cell-cell interaction to affect cell distribution** |  |  | ✓ |
| Transcriptomics | Allow spatial (regional) variation | ✓ | ✓ | ✓ |
|  | Keep gene-gene correlation | ✓ |  | ✓ |
|  | **Allow cell-cell interaction to affect expression – cell distribution association** |  |  | ✓ |
|  | **Allow cell-cell interaction to affect expression – expression association** |  |  | ✓ |

To address this gap, we introduce sCCIgen, a *s*patial resolved transcriptomics *C*ell-*C*ell *I*nteraction *gen*erator, designed to simulate high-fidelity SRT data with a focus on cell-cell interactions. Recognizing the superiority of real-data-based simulators over those relying on preset theoretical models in scRNAseq simulation, sCCIgen utilizes real transcriptomics data for simulation. Unlike existing simulators, it accommodates both single-cell transcriptomics data, such as from scRNAseq/snRNAseq, and single-cell SRT data containing both spatial and transcriptomics information. sCCIgen enhances the strengths of existing simulators by meticulously modeling both the spatial and transcriptomic characteristics of input data and introducing novel cell-cell interaction patterns. This capability bridges a crucial gap in current simulation tools, enabling researchers to delve deeper into the complexities of cellular communication.

## Results

### Overview of sCCIgen

sCCIgen contains the simulation of two major components: the spatial map and gene expression profile **(Fig. 1; Methods)**. With the flexibility to accept input with or without spatial information, sCCIgen either mimics the existing spatial data or generates parameter-guided de novo spatial patterns to simulate the spatial map. When mimicking existing spatial data, sCCIgen accurately estimates the spatial window of the harbored cells, regardless of the slide shape, for both the entire data and separate regions. Within the spatial window, it models the spatial distribution of the cells, allowing for different distributions by cell types and regions, and simulates the spatial map by predicting the spatial coordinates of new cells. When generating spatial data with *de novo* patterns, sCCIgen flexibly simulates different numbers and shapes of spatial regions, and within each region different numbers of cells, cell types, and their composition. It emulates real data by preventing cell overlaps and maintaining balanced cell densities on the slide. Moreover, it enables the manipulation of cell-cell interactions on spatial maps, by attracting or inhibiting cells of the same or different cell types from appearing in the neighborhood.

**Figure 1:**
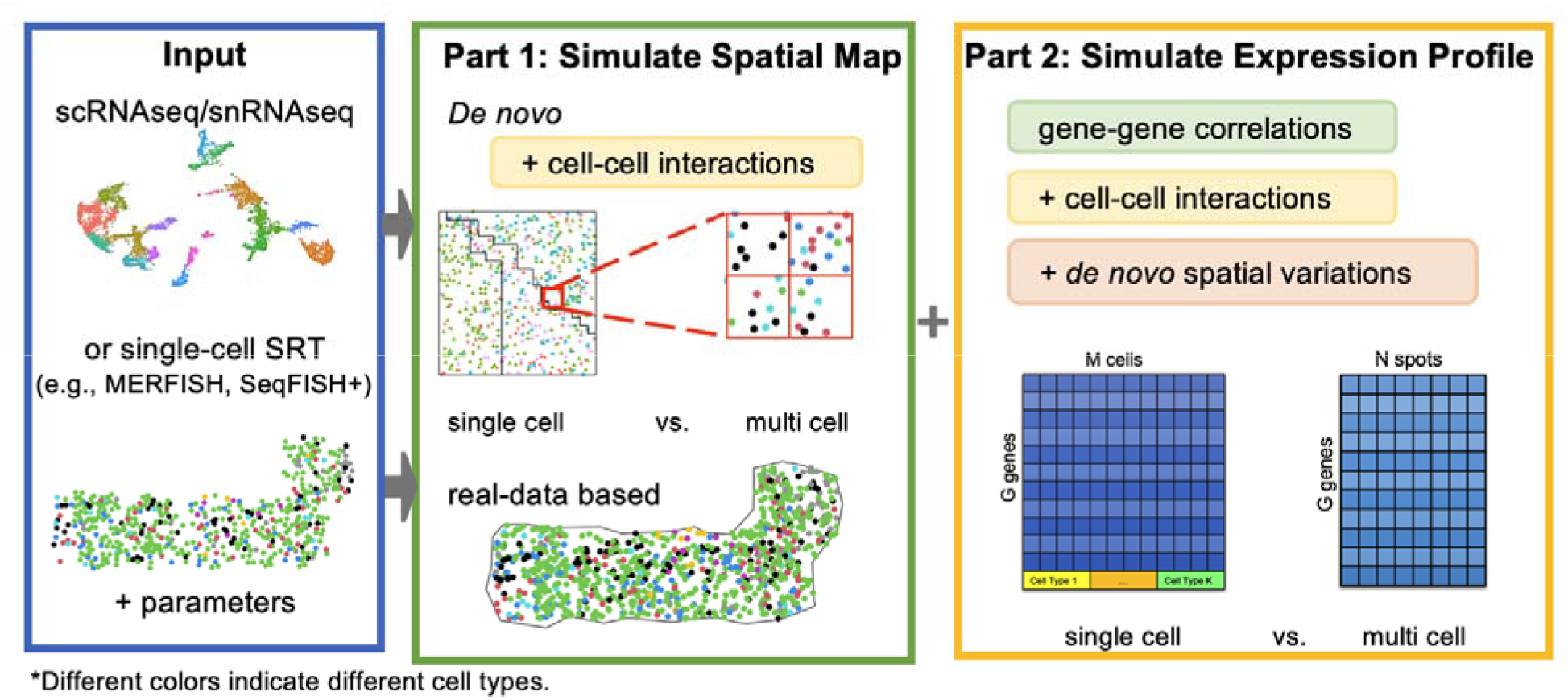
Overview of sCCIgen. sCCIgen operates as a real-data-based simulator, accepting input from single-cell transcriptomics and spatial transcriptomics. It first simulates spatial maps for cells and subsequently generates the expression profile for each cell. The resulting output can be in either single-cell or multi-cell resolution.

Regarding the simulation of the expression profile, sCCIgen relies on the reference transcriptomics data at single-cell resolution. It learns parameters at tissue, cell type, and region levels, such as marginal distributions of genes, gene-gene correlations, and sequencing depths, which can be revised or directly used to simulate expression profiles for cells based on their spatial coordinates. Users can adjust cell-cell interactions on expression by modifying gene expression based on proximity to other cell types (expression-distance association) or by altering gene expression levels based on expression levels of other genes in the neighboring cells (expression-expression association). As a spatial data tool, sCCIgen can introduce spatial expression variations. When combined with cell-cell interactions, sCCIgen generates complex spatial patterns that allow for the evaluation of analytical tools under intricate conditions. Finally, sCCIgen outputs data at single-cell or multi-cell levels, accommodating tools designed for different resolutions.

sCCIgen is available through the widely used programming language R. To facilitate its usage, it also holds a user-friendly interface through Shiny to guide users to generate a well-documented parameter file and use the parameter file to perform reproducible simulations. It also hosts several reference and pre-simulated datasets for users to examine their analytical strategies without performing customized simulations. sCCIgen is publicly available at https://github.com/songxiaoyu/sCCIgen.

### sCCIgen introduces various cell-cell interaction patterns

To facilitate researchers for exploring the dynamic nature of cellular interactions, sCCIgen is designed to introduce cell-cell colocalization, expression-distance association, and expression-expression association. To demonstrate its capacities, we presented SRT datasets **(Fig. 2)** simulated using the snRNAseq data obtained from mammary tissue of a woman with European Ancestry in the Genotype-Tissue Expression (GTEx) project. The reference data included a total of 4751 genes from 5990 cells of six cell types, including epithelial cell, adipocyte, fibroblast, endothelial cell, immune, and others.

**Figure 2.**
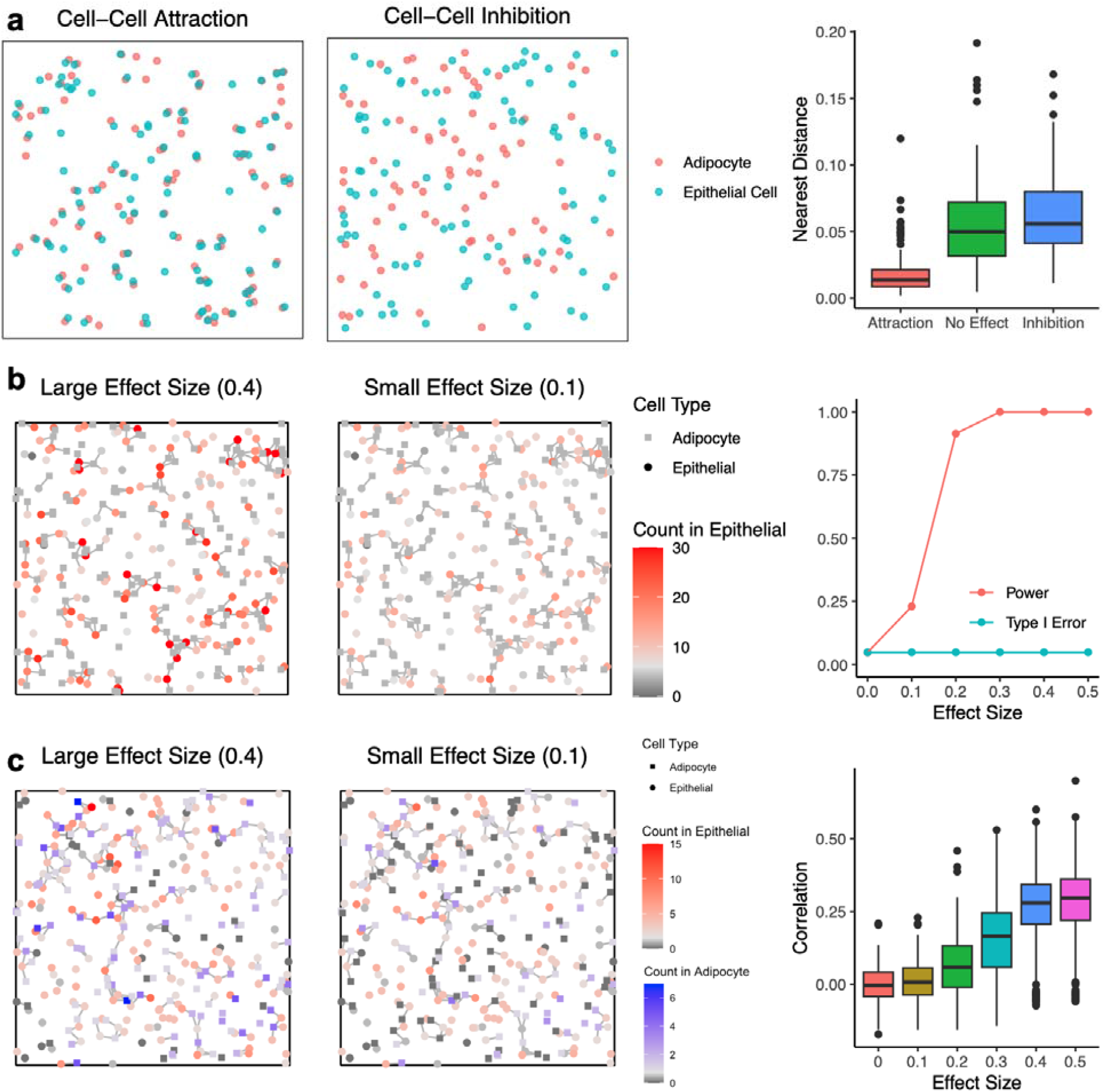
Demonstration of the different *de novo* cell-cell interaction patterns simulated by sCCIgen based on the normal human mammary scRNAseq data. (a) The spatial maps and boxplots of the simulated cells under the cell-cell location attraction/inhibition patterns. The adipocytes and epithelial cells attract (left) or inhibit (middle) each other from locating in the neighbors. The boxplots (right) capture the distance from each adipocyte to its nearest epithelial cell in simulated datasets that have cell-cell attraction, no cell-cell interaction, and cell-cell inhibition. (b) The spatial maps and statistics of the simulated cells under cell-cell expression-distance association patterns. The expression of the demonstrated gene (red-gray) in epithelial cells (circles) is simulated to associate with the existence of nearby adipocytes (squares). An edge represents that two cells are nearby and interact. The effect size is the mean shift on the log-transformed read counts for interacting cells. Type I error and power are reported for identifying the cell-cell interaction changed genes at different effect sizes using a two-sample Wilcoxon test. (c) The spatial maps and statistics of the simulated cells under cell-cell expression-expression association patterns. The expression of the demonstrated gene (red-gray) in epithelial cells (circles) is simulated to associate with the demonstrated gene (blue-gray) in adipocytes (squares). An edge represents that two cells are nearby and interact. The effect size is the mean shift on the log-transformed read counts of both genes in the interacting cells. The boxplots capturing the correlations of these interacting genes are reported for different effect sizes.

To demonstrate the cell-cell colocalization, we first simulated different SRT datasets by alternating the values for the adipocyte-epithelial cell-cell colocalization parameter. The simulated adipocytes and epithelial cells are placed on the spatial map as if they attract **(Fig. 2a; left)** or inhibit **(Fig. 2a; middle)** each other from locating in their neighborhoods. Boxplots capturing cell-cell distance from each adipocyte to its nearest epithelial cell demonstrated their deviations from no cell-cell interactions **(Fig. 2a; right)**. Next, to demonstrate the expression-distance association, we simulated that 10% of the genes in the epithelial cells were positively perturbed by the presence of nearby adipocytes. The expression level of an interacting gene in epithelial cells, at different perturbation effect sizes, together with spatial coordinates with their nearby versus distant adipocytes, were demonstrated **(Fig. 2b; left-middle)**. The larger the effect size, the greater the expression levels were observed in the interacting cells, which were cells with edges to nearby adipocytes, while the expression levels in non-interacting cells with no edges remained the same. Such data could be used for understanding study powers, as well as type I error rates, under different effect sizes for existing analytical tools **(Fig. 2b; right)**. Finally, to demonstrate the expression-expression association, we simulated 10% of genes in adipocytes and the same number of genes in epithelial cells paired with each other and co-expressed in the nearby interacting adipocyte-epithelial cell pairs. The expression levels of an illustrating gene pair in these two cell types, at different effect sizes, were shown **(Fig. 2c; left-middle)**, together with Pearson correlations of these 10% co-expressed gene pairs at different effect sizes **(Fig. 2c; right)**. The larger the effect size, the greater the expression levels for the pair of genes were observed in the interacting cells, while the expression levels remained the same in non-interacting cells.

## sCCIgen facilitates the examination of a complex interplay between spatial variations and cell-cell interactions

sCCIgen builds in features to generate various spatial patterns, facilitating the examination of the complex interplay between spatial variations and cell-cell interactions. It introduces spatial patterns in two ways. First, it allows cell-type composition to vary in different regions, placing cells within distinct microenvironments. Second, it allows certain genes in some cell types to be activated or suppressed in particular spatial regions, such as in response to pathological triggers. Even when cells respond similarly to signals from nearby cells, these spatial patterns can influence their behaviors. By building in features to simulate both cell-cell interactions and spatial variations, sCCIgen provides a realistic depiction of cellular behaviors under their intricate interplay in real-world data.

To demonstrate the interplay between cell-type composition variation and their interaction, we simulated a dataset with three spatial regions of diverse cell-type composition (**Fig. 3a**). The reference data was again the snRNAseq data from mammary tissue. Across all three regions, the epithelial cells were generated to interact with their neighboring immune cells if they were located close to each other, presenting an edge in **Fig. 3b**. However, the frequency of interactions varied according to the differing densities of epithelial and immune cells, with interactions most frequent in the middle region where the two cell types intermixed the most (**Fig. 3b**). We simulated that these interactions perturbed 10% of randomly selected genes, termed interaction-change genes (ICGs), which on average increased the log-transformed count level by per interaction pair (edge). The normalized averaged expression levels of these ICGs, as well as the epithelial signature genes (SGs), in each epithelial and non-epithelial cell are presented (**Fig. 3c)**. The epithelial SGs were identified in the reference snRNAseq data from one-sided differential expression analysis comparing epithelial cells with all other cell types combined at 10% false discovery rate (FDR). Clearly, ICGs were higher in simulated cells in the middle region than others, epithelial SGs were higher in simulated epithelial cells than other cells, and their overlapping genes were especially elevated in the simulated epithelial cells of the middle region. The multi-cell resolution data presented the same pattern with enhanced noises (**Fig. 3d)**. Spatial variable gene (SVG) analysis under cell-cell interactions versus no interactions demonstrated that cell-cell interaction drove some genes to become SVG while reducing the p-value of other genes (**Fig. 3e)**, underscoring the importance of considering cell-cell interactions on the analysis and interpretation of spatial variable genes.

**Figure 3.**
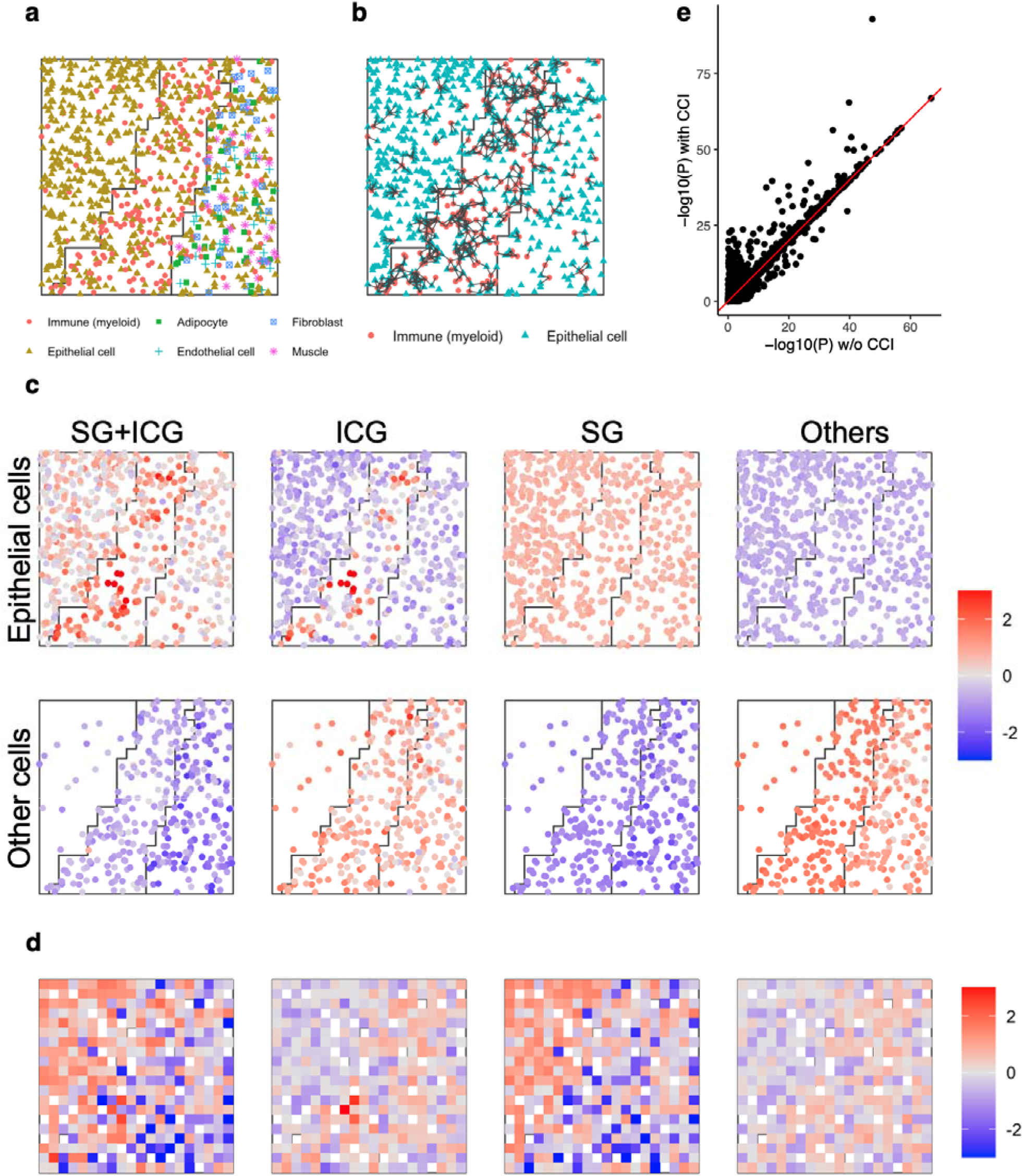
Demonstration of the compounding effects of cell-cell interaction and spatial variation on cell-type composition, using sCCIgen simulated data based on the normal human mammary scRNAseq. (a) The spatial map of the simulated cells of six cell types. (b) Illustration of cell-cell interaction between epithelial and immune cells on the spatial map. An edge represents that two cells are nearby and interact. (c) Averaged expression levels in epithelial and other cell types for simulated genes divided into four exclusive groups: epithelial signature gene (SG) + interaction-changed genes (ICG), ICG only, SG only, and others. (d) Averaged expression levels at the multi-cell resolution for the four groups of simulated genes. (e) P-values from spatial variation analysis for all genes in simulated data versus w/o epithelial-immune cell-cell interaction.

We can similarly evaluate the interplay between cell-cell interactions and the spatial variations introduced by perturbed expressions in different regions of the same cell type. We simulated the same three spatial regions as before, but this time different cell types were positioned in the same way in these regions (**Fig. 4a**). As a result, the simulated epithelial-immune interactions occurred in similar frequencies in different regions. (**Fig. 4b)**. In addition to the interactions, we also introduced spatial variation by increasing the expression levels for 20% of genes in epithelial cells in the middle region, designating this special case of spatial variable genes as regional variable genes (RVGs). The normalized averaged expression levels of these RVGs and ICGs, in each epithelial and non-epithelial cell are presented (**Fig. 4c**). It illustrated that RVGs had higher expression levels in the middle region than other regions in the simulated epithelial cells and ICGs exhibited clustered elevated expression levels but did not show any regional patterns, which was consistent with the multi-cell resolution output data (**Fig. 4d)**. Spatial variable gene analysis prioritized different genes under the existence of these cell-cell interactions versus no interactions (**Fig. 4e)**, again suggesting that cell-cell interactions have a significant impact on the identification of spatial variable genes.

**Figure 4.**
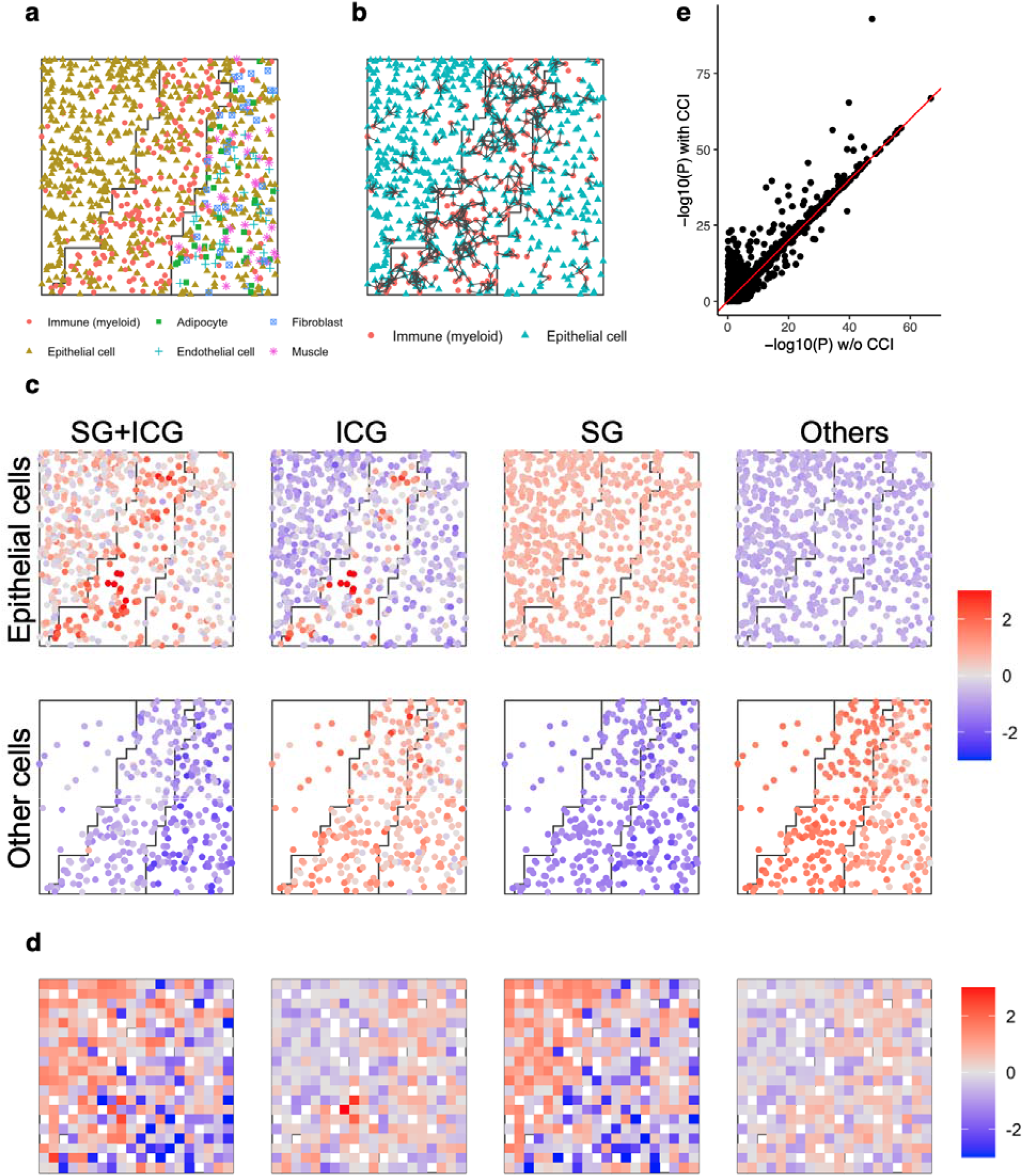
Demonstration of the compounding effects of cell-cell interaction and regional differential expression within the same cell type, using sCCIgen simulated data based on the normal human mammary scRNAseq. (a) The spatial map of the simulated cells of six cell types. (b) Illustration of cell-cell interaction between epithelial and immune cells on the spatial map. An edge represents that two cells are nearby and interact. (c) Averaged expression levels in epithelial and other cell types for simulated genes divided into four exclusive groups: regional variation gene (RVG) + interaction-changed genes (ICG), ICG only, RVG only, and others. (d) Averaged expression levels at the multi-cell resolution for the four groups of simulated genes. (e) P-values from spatial variation analysis for all genes in simulated data versus w/o epithelial-immune cell-cell interaction.

### sCCIgen mimics the spatial and transcriptomic features of the reference data

sCCIgen designs several critical features to closely approximate the various spatial and transcriptomic characteristics of the reference data. To demonstrate these features, we simulated 1000 cells using a reference SRT dataset from a mouse cortex, profiled using SeqFISH+ technology^17^ (**Figure 5**). Simulation utilized 2,000 highly variable genes from 511 cells of six cell types, including excitatory neuron, interneuron, astrocyte, endothelial, oligodendrocyte, and microglia.

**Figure 5.**
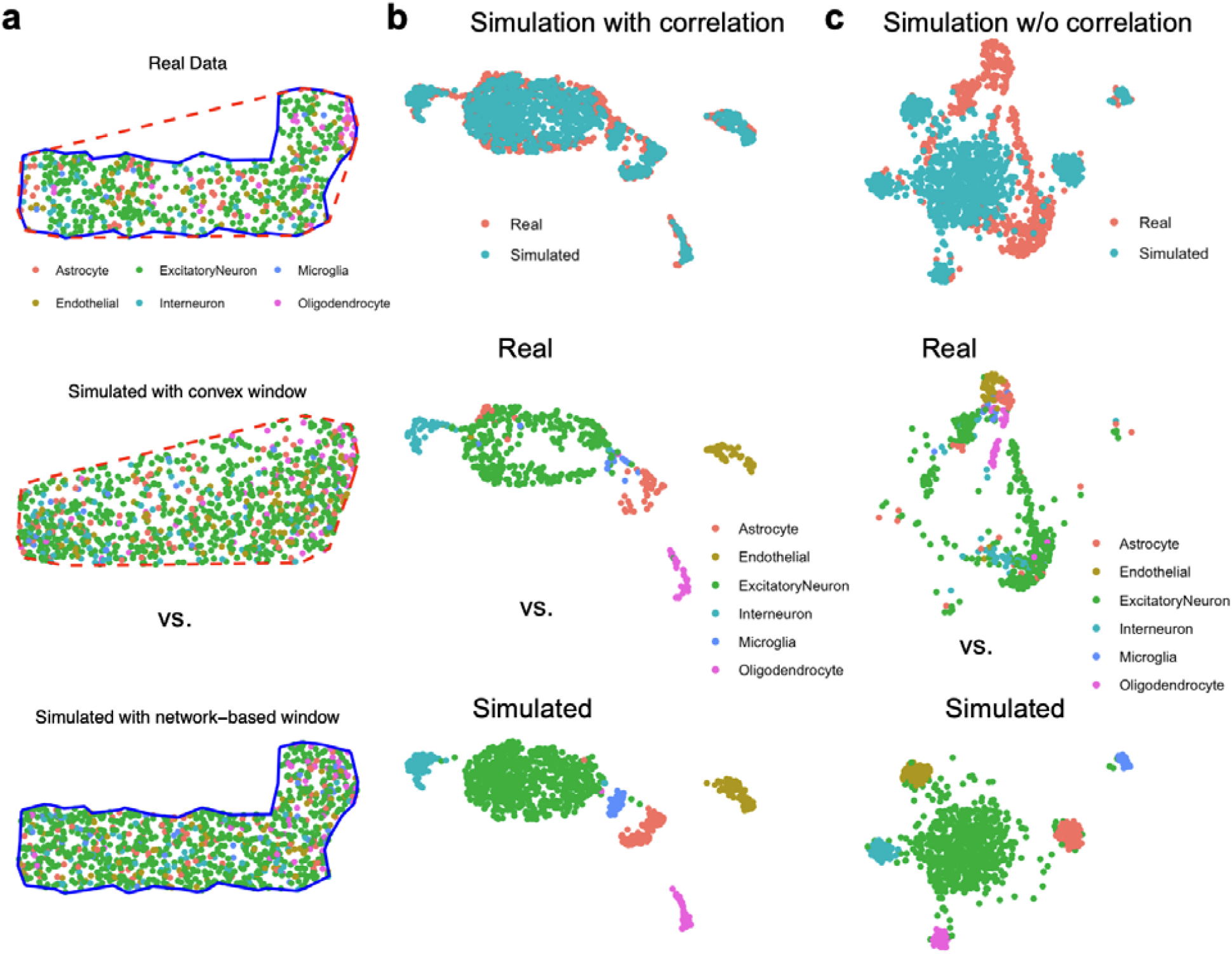
sCCIgen simulates data that highly mimics the spatial and transcriptomic features of reference data. (a) Spatial maps of real data, simulated data using conventional window estimation, and simulated data using sCCIgen-developed network-based window estimation. (b) UMAP of real data compared to simulated data capturing gene-gene correlation. (c) UMAP of real data compared to simulated data without capturing gene-gene correlation.

First, sCCIgen enhances the generation of spatial maps. It develops a network-based window estimation algorithm to capture the complex geometries of the input data (Methods). An estimation of the spatial window helps prevent cells from being placed outside the areas covered by the reference data. The algorithm effectively captures the complex shapes of the input data, resulting in improved cell placement compared to using simple shapes like rectangles or convex polygons as used in existing simulators (**Fig. 5a**). sCCIgen also includes features to ensure there are no overlapping cells on the spatial maps and promote an even cell distribution, preventing large empty spaces within the window. These features make the spatial map of simulated data highly mimic the real reference data. Second, sCCIgen carefully models the transcriptomics features of the data, including gene-gene correlations, resulting in simulated data that closely resembles real reference data (**Fig. 5b**). This approach provides a stark contrast to simulations that overlook gene-gene correlations (**Fig. 5c**).

## Discussion

The advent of SRT technologies has provided researchers with a powerful tool to dissect the intricate landscape of cell-cell interactions within the native tissue context. In this manuscript, we introduced sCCIgen, a novel SRT data simulator designed to address the critical need for simulating high-fidelity SRT data with cell-cell interactions. sCCIgen not only integrates and enhances the strengths of existing simulators in replicating the spatial and transcriptomic features of reference data, but it also simulates diverse cell-cell interaction patterns and *de novo* spatial patterns. It facilitates simulations based on transcriptomics without spatial input and enables the examination of the interplay between spatial variations and cell-cell interactions. sCCIgen offers an R package with a user-friendly interface that includes detailed instructions, tutorials, and examples, ensuring ease of use for researchers with varying levels of computational expertise.

sCCIgen can be naturally used for benchmark analyses of various cell-cell interaction tools. It can also be used to explore study design and sample size considerations to plan experiments under different effect sizes. Additionally, sCCIgen can be employed to evaluate the impact of cell-cell interactions on diverse spatial analyses. Our analysis indicates that cell-cell interactions can influence the detection of spatial variable genes, implying potential effects on spatial clustering, cell-type deconvolution, and cell trajectory analysis. Since few tools consider cell-cell interactions during development, sCCIgen can assess their performance in light of these ubiquitous signals, paving the way for future studies to refine their analytical methods and better understand the complex dynamics of spatially resolved transcriptomic data.

Since sCCIgen is designed with a focus on cell-cell interactions, it is limited to input data at the cellular resolution. Future studies could expand its capabilities by integrating sCCIgen with emerging cell-cell interaction tools that operate at multi-cell or subcellular resolutions, allowing it to process information at different scales. Additionally, sCCIgen is primarily centered around transcriptomics data, while spatial genomics and proteomics data are becoming increasingly available. The count model built into sCCIgen may not be directly applicable to these additional layers of datasets. Extending the statistical models to better describe these additional omics data would enhance sCCIgen’s utility for exploring complex spatial biology.

## Methods

### sCCIgen Step 1: Simulate the spatial map for cells

sCCIgen leverages as much spatial information from the input reference data as possible for simulation. When the input data lacks spatial information, such as from snRNAseq, sCCIgen guides users to specify parameters and use these parameters to simulate the spatial maps for cells. Alternatively, sCCIgen learns parameters from the data for simulation.

### Simulation without spatial reference

For simulations lacking spatial reference, users define parameters to specify attributes. If multiple spatial regions are specified, sCCIgen develops a random-walk-based algorithm to randomly segregate the spatial window into different regions. Within each region, it generates cell locations based on user-specified cell numbers and cell-type proportions. Uniquely, sCCIgen allows users to specify spatial attraction and inhibition patterns of cells, mimicking cell-cell interactions impacting their spatial distributions. It also includes features to smooth cell densities and remove cell overlaps, mirroring real tissue scenarios.

### Segregate spatial regions

We developed a random-walk-based algorithm to generate randomly connected spatial regions within a window. When users specify K regions, where K>1, sCCIgen first partitions the window into B by B tiny rectangular pixels arranged in a grid formation, where B (e.g. B=20) is a nuisance parameter relevant to the smoothness of region boundaries. Next, sCCIgen identifies neighbors of these rectangular pixels that share a common edge. Pixels with less than four neighbors are on the border of the window, and K of them are randomly selected as the first pixel of the regions. Then for each region, sCCIgen finds all neighbors of its pixel(s) that have not been assigned to any regions and randomly selects one as a random walk for this region to grow. sCCIgen runs this random walk one step in a region and then goes to the next region until all pixels have been assigned to the regions. This random-walk-based algorithm partitions the spatial windows into K randomly connected regions with similar sizes.

### Simulate cell coordinates within a region

Within each region, sCCIgen generates spatial coordinates of cells that satisfy the user-specified parameters. In order to achieve the targeted cell numbers for different cell types and regions while satisfying the user-specified spatial patterns, such as cell-cell spatial attraction or inhibition, cell distribution evenness, and cell nonoverlap, sCCIgen first inflates the cell numbers to create a pool of cells to choose from. The inflation is linearly dependent on the absolute strength of cell-cell inhibition/attraction and to the power of (1 + cell distribution evenness parameter). The stronger cell-cell attraction/inhibition and the greater cell distribution evenness users specify, the bigger the inflated cell numbers will be. Next, sCCIgen generates the spatial coordinates for these inflated numbers of cells under the uniform distribution. Then, sCCIgen implements two selection mechanisms to harvest the targeted number of cells from the pool. When cells are located within the minimally allowed distance indicating they are overlapping, all but one cell are randomly deleted. In the remaining cells, it calculates the selection probability for cells in each cell type based on (1) attraction and inhibition of the same cell type, (2) attraction and inhibition of different cell types, and (3) even distribution requirement of cells. Specifically, for each cell in cell type, the logit form of selection probability is

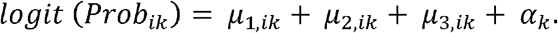

The parameter *µ*_1,*ik*_ controls the impact of cell-cell interaction from the neighboring cells of the same cell type. It is calculated as the product of the density of cells in the same cell type in the neighborhood and the cell-cell inhibition/attraction strength specified by users. When cells inhibit each other from occurring in the neighborhood (strength<0), *µ*_1,*ik*_ < 0, meaning neighboring cells reduce the selection probability of the cell. When cells attract each other from occurring in the neighborhood (strength>0), *µ*_1,*ik*_ >0, meaning neighboring cells increase the selection probability of the cell. Similarly, *µ*_1,*ik*_ controls the impact of cell-cell interaction from the neighboring cells of the different cell types, calculated as the product of the density of cells in the different cell types in the neighborhood and the user-specified cell-cell inhibition/attraction strength. The parameter *µ*_1,*ik*_ controls the cell distribution evenness, calculated as the product of the density of all cells in the neighborhood and the cell distribution evenness parameter. Finally, *α*_*k*_ is a nuisance parameter calculated by sCCIgen to ensure the selection probability across all cells matches the ratio between the target and existing cell number in the pool for this cell type. Based on this selection probability, cells are randomly selected, exhibiting the user-specified spatial patterns. The cell features including their cell types, spatial coordinates, and regions are kept as an output to users.

### Simulation with spatial reference

In some datasets such as MERFISH and SeqFISH+, spatial information is available including spatial coordinates (x, y) and sometimes regions. sCCIgen can estimate the spatial window of these cells to ensure simulation within the same area of input, model the spatial patterns of the cells, and predict the spatial map for cells based on the models. If input data covers multiple regions, sCCIgen performs region-specific simulation and integrates the spatial map for cells across all regions. Alternatively, if users would like to use the existing spatial maps of cells, they can skip this step to simulate the expression profiles of cells.

### Estimate the spatial window

sCCIgen develops a novel network-based algorithm to accurately estimate the spatial window for input spatial data with complex shapes.

Specifically, it first builds the Delaunay triangulation^18^ network using the cell spatial coordinates. Then, it identifies the outlier edges in the network, such as those three standard deviations (SD) longer than the averaged network length (e.g. > mean+3SD). Next, it summarizes an outlier-excluded network and finally extracts the outer frame of the network as the window of the ST data. In addition to providing this accurate widow estimation algorithm, sCCIgen also allows for a fast approximation using simple functions such as drawing a “rectangular” or “convex” polygon as in existing simulators. It also allows users to select a function to split the data into 2-5 sections, estimate the “convex” polygons in each section, and merge the polygons to estimate the window, which balances between accuracy and complexity.

### Simulate the cell spatial locations

Within the window, sCCIgen fits a parametric Poisson point process model^19^ to model the logarithm density of cells as a function of the cell types, the polynomial form of spatial coordinates (x, y, x^2, y^2, xy), and their interactions. Next, the spatial maps of the user-specified numbers of cells are generated by predicting the spatial coordinates of new cells using the fitted Poisson point process model.

### sCCIgen Step 2: Simulate expression profiles of the cells

#### Generate the initial read counts based on the input single-cell expression data

Leveraging the existing scRNAseq simulator scDesign2^20^, we fit the marginal distribution of each gene using (zero-inflated) Poisson or negative binomial distributions for each cell type. If gene-gene correlations are specified, we also estimate the correlation for each cell type using a Gaussian copula. This model fitting procedure can be region-specific, as determined by the input data and user-specified parameters. Then, for simulated cells with spatial maps in Step 1, we simulate their gene expression profiles by predicting the counts based on the model, using user-specified sequencing depth. These predicted counts are cell-type specific and can be region-specific as determined by users’ specifications.

#### Add the impact of cell-cell interaction on the expression-distance association

Based on the initial read counts generated from the reference data, sCCIgen allows users to alter the expression levels of some genes in some cells based on the presence of other cells in the local microenvironment, thereby building their expression-distance associations. Users need to specify parameters such as the interaction region(s), perturbed cell type(s), neighbor cell type(s), interacting distance threshold(s), perturbed gene(s), and the mean and standard deviation of effect size(s). These perturbed genes can be provided by users, such as from the estimation of existing data using methods that study cell-cell interactions^21^, or can be randomly generated by specifying the percentage of perturbed genes. The effect sizes are applied to the log-transformed counts, and thus negative perturbations on genes already with zero counts will not change expression levels in these cells.

#### Add in the impact of cell-cell interaction on the expression-expression association

Similarly, sCCIgen allows users to alter the expression levels of some genes in some cells based on the expression levels of some other genes in some other cells in the local microenvironment, thereby building their expression-expression associations. Users need to specify parameters such as the interaction region(s), perturbed cell type(s), neighbor cell type(s), interacting distance threshold(s), perturbed gene pair(s), whether the perturbation is bidirectional, and the mean and standard deviation of effect size(s) of the perturbation. These perturbed gene pairs can be estimated using existing data or be randomly generated. The effect sizes are applied to the log-transformed counts.

#### Add in the regional variation

To mimic the spatial variation of gene expression, sCCIgen requires users to specify spatial regions of the input data or the number of regions in the simulated data and can add regional variations on the expression profiles. Users need to specify parameters such as the perturbed region(s), cell type(s) and gene(s), and the mean and standard deviation of effect size(s) of the perturbation. These perturbed genes can be provided by users, such as by estimating the spatial variable genes in the existing data^21^ or randomly generated with effect sizes applied to the log-transformed counts.

#### Read counts

Multiple patterns can be added simultaneously, and their cumulative effects on the log count scale will be put on the genes. After incorporating these associations, users can specify parameters to limit the extreme values of the counts and neutralize their impact on sequencing depth.

#### sCCIgen multi-cell simulation

When generating multi-cell SRT data, sCCIgen cuts the window into squares of equal sizes. The number of squares are specified by users and suggested to cover ∼0-40 cells mimicking a predetermined SRT spot. The spatial coordinates are the centers of the squares, and the expression levels are summed for all cells in the squares.

#### User-friendly interface

To facilitate the usage of sCCIgen, we developed an interactive Shiny application using the packages Shiny (version 1.8.1)^22^ and shinyFiles (version 0.9.3)^23^. The interactive application is contained in the function run_interactive_sCCIgen() which launches a local server-free shiny interface. The application contains three main options: 1) download a previously simulated dataset, 2) create a parameter file, or 3) run a simulation. In addition to downloading pre-simulated datasets, the app provides the option of downloading test and real datasets that will be used as reference for running the simulation. Alternatively, the user can select their own local expression and metadata files as the reference data. When creating a parameter file, the application shows multiple interactive questions that will lead the user through a series of customized options to select the proper parameters and values depending on the dataset of reference used. Finally, we integrated sCCIgen with the Giotto package (version 4.0.6)^24^ by providing the option of creating and saving a Giotto object that contains the simulated dataset, in addition to the default output files. The integration with Giotto will facilitate running downstream spatial analysis with this package.

#### Three datasets are hosted by sCCIgen

sCCIgen provides three real reference datasets, from snRNAseq, SeqFISH, and MERFISH, for users to explore and perform simulation. The snRNAseq data was from the mammary tissue of one European Ancestry woman ‘‘GTEX-1R9PN’’ downloaded from the Genotype-Tissue Expression (GTEx) project. The datasets included the expression profile for a total of 4751 genes from 5990 cells of six cell types: epithelial cell (n=2292), adipocyte (n=1721), fibroblast (n=804), endothelial cell (n=737), immune (n=301), and other (n=135). The SeqFISH+ reference dataset was obtained from the cortex of a 23-day-old male mouse (C57BL/6J). The dataset included the spatial maps of 511 cells and expression levels for 10,000 genes. These cells were categorized by the original study into six major cell types: excitatory neuron (n=325), interneuron (n=42), astrocyte (n=54), endothelial (n=45), oligodendrocyte (n=29), and microglia (n=16). The MERFISH dataset downloaded from Vizgen was from an ovarian cancer sample including a total of 550 genes from 355,633 cells. We annotated six cell types in the data preparation step, including tumor (epithelial) cells, adipocytes, T cells, endothelial cells, macrophages, and others.

## Acknowledgments

This work was supported by NIH grants R03AG075567, RF1MH133703, and RF1 MH128970, and by the Duke-NUS Signature Research Programme funded by the Ministry of Health, Singapore.

